# Inhibited oligodendrogenesis, but not repeated mild traumatic brain injury, impairs attention in adult mice

**DOI:** 10.1101/2025.09.01.673570

**Authors:** Lisa M. Gazdzinski, Jordan Mak, Paul J Fletcher, Anne L. Wheeler

## Abstract

Attention problems are among the most common long-lasting cognitive symptoms of mild traumatic brain injury (mTBI) and as attention is fundamental to many aspects of cognition, the effects of attentional impairment can be broad. The brain’s white matter is particularly vulnerable to damage during mTBI. Damage to oligodendrocytes and myelin contributes to cognitive deficits following injury and myelin plasticity is a potential mechanism for functional recovery. The aim of this work was to assess attentional impairment following mTBI in mice and evaluate the role of newly generated oligodendrocytes in recovery. This study used the Myrf conditional knockout mouse model, in which the Myrf gene, required for oligodendrocyte precursor cell (OPC) differentiation into mature myelinating oligodendrocytes, is deleted from OPCs following tamoxifen injection, thereby halting oligodendrogenesis. Mice were trained on the 5-Choice Serial Reaction Time (5-CSRT) task before receiving tamoxifen followed by three mTBI or sham procedures. Attention was probed on the 5-CSRT with decreasing stimulus durations at four time points following injury out to 12 weeks. While no attentional impairment was observed in mice with mTBI, OPC-MyrfKO mice showed lower choice accuracy (β = −2.85%, p = 0.03) and more omitted trials (β = 4.09%, p = 0.02) across injury groups, time points and stimulus durations, suggesting that active oligodendrogenesis is required for sustained attention. These results suggest that this mouse model of mTBI was not severe enough to impact attention as measured by the 5-CSRT task, however myelin plasticity in adulthood may contribute to attention and complex task performance.

**Significance Statement:** Attention deficits are common after mild traumatic brain injury (mTBI), yet therapeutic targets remain unclear. This study investigated the role of adult myelin plasticity in cognitive recovery by using a mouse model that blocks new myelin formation. While mild injury alone did not impair attention, mice unable to generate new myelin showed reduced response accuracy. These findings suggest that adult oligodendrogenesis supports attention. Understanding how myelin plasticity influences behaviour could reveal new targets for enhancing recovery after brain injury, highlighting the importance of supporting myelin health in therapeutic strategies.

## 1 Introduction

Mild traumatic brain injury (mTBI), also often called concussion, is very common, with an estimated 1 in 4 people being affected in their lifetime (Daugherty et al., 2020). Repeated injuries are common, particularly among contact sport athletes (Glaser et al., 2024; Yates et al., 2025) and in people who experience intimate partner violence (Smirl et al., 2019). While most recover fully within a few days to a couple of weeks, a significant proportion (∼30%) of people experience prolonged symptoms that can last months or years after the injury (Cancelliere et al., 2023). A recent systematic review listed impaired attention among the most common cognitive deficits following concussion from the acute through to the long-term period (Collins et al., 2023). Attention deficits have also been reported in pre-clinical TBI models. Impaired performance on the 5-choice serial reaction time (5-CSRT) task has been observed in juvenile and adult rats following repeated mTBI (Mychasiuk et al., 2015; Xu et al., 2021; Vonder Haar et al., 2022), and a transient post-mTBI deficit has been reported in mice in the object-based attention test (Zhang et al., 2021) and in the Y-maze spontaneous alternation task (Gazdzinski et al., 2020), which is considered a test of working memory, but also requires attentional processes for performance. As attention is fundamental to many other aspects of cognition, attentional impairment can impact many domains of daily functioning for people after mTBI, thus therapeutic targets are needed.

White matter, which contains bundles of axons responsible for communication across the brain, is particularly vulnerable to damage from TBI (Armstrong et al., 2016). Axons may be directly damaged by stretching and shearing forces imparted on them by the acceleration and deceleration of the head during the injuring event itself (Johnson et al., 2013), but if the injury is mild, it does not necessarily lead to axonal loss and cell death. Instead, mild trauma can result in damaged axons dispersed among intact ones within major white matter tracts (Povlishock, 1993; Sullivan et al., 2013). Many large caliber axons are surrounded by myelin sheaths. Myelin is formed from multiple layers of extended cell membrane from oligodendrocytes, with a single oligodendrocyte contributing myelin to several axons. In the damaged white matter environment that results from mTBI, oxidative stress, glutamate excitotoxicity, and inflammation can lead to loss of oligodendrocytes, demyelination of intact axons, and changes to myelin structure (Lotocki et al., 2011; Shi et al., 2016). Given myelin’s role in supporting and tuning neural circuit function, myelin damage may be a significant factor underlying the cognitive impairments following mTBI (Mierzwa et al., 2015). Fortunately, evidence for myelin plasticity and repair after mTBI has been found in both human imaging studies, which have shown evolution and in some cases normalization of myelin water fraction and diffusion MRI measures over time (Weber et al., 2018; Lindsey et al., 2023), and in animal models of mTBI, in which increased oligodendrocyte precursor cell (OPC) proliferation and evidence of remyelination have been reported (Sullivan et al., 2013; Mierzwa et al., 2015; Nonaka et al., 2021). Thus, myelin plasticity is a potential mechanism for functional recovery from mTBI and understanding its role is important for assessing it as a therapeutic target.

The objectives of the current study were to assess attentional impairment and recovery following mTBI in mice and to investigate the contribution of new myelin to recovery. The study was conducted in Myrf conditional knockout (Myrf-cKO) mice, in which the myelin regulatory factor (Myrf) gene, a transcription factor required for OPC differentiation, is deleted from OPCs in a time-controlled manner via tamoxifen administration (McKenzie et al., 2014; Duncan et al., 2018; Pan et al., 2020; Steadman et al., 2020; Xin et al., 2024). Mice were trained on the 5-CSRT task, a pre-clinical adaptation of the continuous performance test commonly used to assess attention in humans (Mar et al., 2013; Fizet et al., 2016). After training, oligodendrogenesis was halted in Cre+ Myrf-cKO mice (OPC-MyrfKO) before the mice underwent repeated mTBI or sham procedures. Attention was assessed by decreasing the stimulus duration on the 5-CSRT task at four post-mTBI timepoints out to 12 weeks. While attentional impairment was not observed following mTBI in this study, mice with disrupted oligodendrogenesis omitted more trials, had lower choice accuracy, and were slower to collect the reward across injury groups, stimulus durations and time points compared to Cre-controls (OPC-MyrfKO-CTRL), suggesting that active myelination is required for sustained attention and engagement in the task.

## 2 Materials & Methods

### 2.1 Mice

All animal procedures were performed in accordance with the [Author University] animal care committee’s regulations. Myrf-cKO mice were generated by first crossing Myrf^*flox/flox*^ mice (The Jackson Laboratory #01067, in-house colony) with NG2-CreER^TM^ mice (The Jackson Laboratory #8538, in-house colony) to generate Myrf^*flox/+*^:NG2-CreER^TM^ mice, which were backcrossed with Myrf^*flox/flox*^ mice to yield mice that were homozygous for the floxed Myrf allele (Myrf^*flox/flox*^) and either hemizygous (Cre+) or non-carrier (Cre-) for the NG2-CreER^TM^ gene. Approximately equal numbers of male and female mice of each Cre genotype were included in each injury group (N = 76; 18-20/genotype/injury group), and mice were housed in sex-separated cages in groups of 2-5. Pre-training began when mice were 8-11 weeks old. Training and testing were performed during the latter half of the light phase of the 12-hour light/dark cycle (ZT6-ZT12). A study timeline is shown in **Figure 1**.

**Figure 1.**
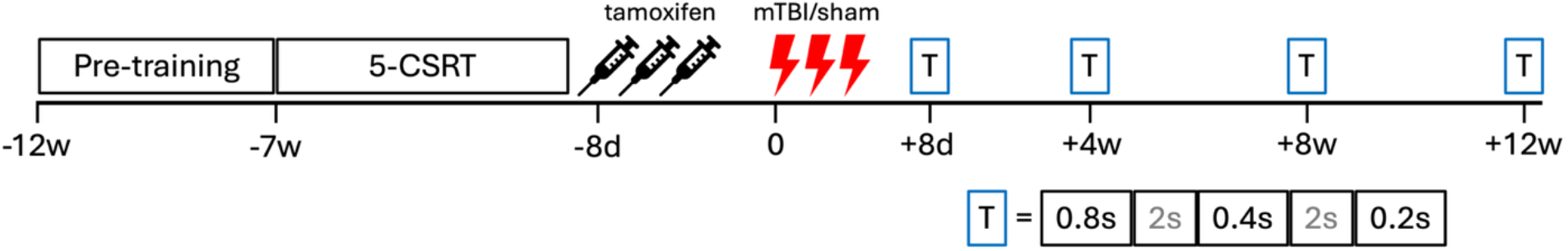
Study timeline. Mice underwent pre-training followed by 5-CSRT training starting about 12 weeks before mTBI. Tamoxifen was given 8 days before the first mTBI procedure to halt oligodendrogenesis in Cre+ Myrf-cKO mice (OPC-MyrfKO). Testing (T in blue box) was performed at 8 days, 4 weeks, 8 weeks and 12 weeks post-mTBI. At each testing timepoint, the probe trial stimulus duration decreased from 0.8 s to 0.4 s to 0.2 s with rebaselining sessions with the 2-s stimulus duration between probe trials.

### 2.2 Tamoxifen administration

Tamoxifen (T5648 Sigma-Aldrich, 180 mg/kg dissolved at 30 mg/ml in 10% ethanol in sunflower oil (Sigma-Aldrich, Canada)) was administered to all mice via daily intraperitoneal injections for three days, which activated Cre recombinase in Cre+ mice, thereby deleting the Myrf gene in NG2-expressing cells in these mice (OPC-MyrfKO). The mice were given supplemental food (DietGel 76A, Clear H2O, USA) beginning immediately after the final task session before tamoxifen and until 24 hrs after the last tamoxifen injection to decrease morbidity associated with the drug.

### 2.3 5-Choice Serial Reaction Time (5-CSRT) Task

Mice were trained in 16 Bussey-Saksida touchscreen operant chambers (Campden Instruments, USA). One wall of the chamber consisted of a touchscreen on which an illuminated square stimulus was presented in one of five locations. The opposite wall of the chamber contained a reward port, in which strawberry milkshake (Neilson, Canada) was delivered following a correct response. Data from one of the chambers (3 mice) was excluded from all analysis because there was an intermittent malfunction, making the data from this chamber unreliable.

#### Food restriction

The mice were food restricted beginning at 6-9 weeks of age and for the duration of the study to maintain a target weight of 80-85% of their free-feeding weight. If food restriction began prior to 8 weeks of age, the target weight was increased by 1 g/week to account for developmental growth. Habituation and training began after 1-2 weeks of food restriction.

#### Habituation and pre-training

The mice were habituated to the chambers and the milkshake over two days and then progressed through several stages of pre-training over the following 2-4 weeks in which they learned to associate touching the stimulus with the delivery of milkshake reward. Details about the stages are provided in the supplemental material.

#### Task-specific training

For this stage of the training, mice were required to initiate each trial by poking their head into the reward port, after which there was a 5-s delay before a stimulus would be presented on the screen. The mice were required to touch the stimulus window either while it was illuminated or during the 5-s limited hold period that followed to receive reward. An incorrect response, an omitted trial, or a touch during the 5 s before stimulus presentation, would end the trial and trigger the chamber light to come on and a buzzer to sound for 5 s. The stimulus duration was progressively reduced from 8 s to 4 s to 2 s. Mice were required to complete 40 correct trials within 60 minutes with an accuracy (# correct/(# correct + # incorrect + # omitted)x100% of at least 70% and fewer than 20% omitted trials on 3 out of 4 consecutive days to progress to the next stimulus duration. This stage of training lasted on average 5-6 weeks with the mice being trained 6 days/week. Mice that completed training earlier than others in their cohort were given weekly reminder sessions with the 2-s stimulus duration.

#### Testing

At 8 days, 4 weeks, 8 weeks, and 12 weeks after the first mTBI procedure, the mice were tested on the task with stimulus durations of 0.8 s, 0.4 s, and 0.2 s. Prior to each of these probe trial series, the mice had three rebaselining sessions with the 2-s stimulus duration. Single rebaselining sessions were also done between each of the probe trials. Weekly reminder sessions with the 2-s stimulus duration were done between post-mTBI timepoints.

#### Measures

The number of correct, incorrect, and omitted trials were counted and from this, choice accuracy (# correct/(# correct + incorrect)x100%) and % omitted trials were calculated. Correct response, incorrect response, and reward collection latencies were also recorded.

### 2.4 Repeated mild TBI

Following tamoxifen administration, the mice remained in their home cage for two days and then performed three rebaselining sessions with the 2-s stimulus duration. The average accuracy from these three sessions was considered when assigning the mice to the mTBI or sham groups such that the injury groups were balanced for task performance as well as genotype (Cre status) and sex. The mice were then subjected to three mild TBI or sham procedures spaced 24 hrs apart. Under isoflurane anaesthesia (4% for induction; 1-2% for maintenance), the hair on the scalp was removed using depilatory cream (Nair, Church & Dwight Co., Inc., USA) and the mouse was positioned in a stereotaxic frame (Stoelting Co., USA) using zygomatic arch cuffs. An electromagnetic controlled cortical impact device (Impact One, Leica Biosystems, USA) was used to deliver an impact to the intact scalp centred at bregma with a 5-mm metal tip to a depth of 1.2 mm at a velocity of 2 m/s and with a dwell time of 0.2 s. Following the impact, the mouse was allowed to breathe oxygen for 1 min and was then transferred to a warm cage to recover from anaesthesia before being returned to its home cage. Subcutaneous saline was provided before each procedure to maintain hydration and sustained-release buprenorphine (1.2 mg/kg) was provided before the first and third procedures to provide analgesia. The mice were given supplemental food following each mTBI procedure, as during tamoxifen administration.

### 2.5 Statistical Analyses

Group comparisons in choice accuracy, % omissions, and correct response and reward collection latencies were made in the probe trial data (0.8 s, 0.4 s, and 0.2 s stimulus durations) using linear mixed effects models that included covariates for injury group, genotype, sex, stimulus duration, and timepoint, and random intercepts for mouse and cohort to account for repeated testing and correlations among mice tested together. The Satterthwaite approximation was used to estimate degrees of freedom for fixed effects. Interaction terms to test for injury group and genotype effects on change in measures over time were initially included, but as no significant interactions were observed, the interaction terms were not included in the final models. Statistical analyses were performed in R (4.4.0; RStudio 2024.09.1+394 for macOS) (R Core Team, 2022).

## 3 Results

The mice learned the 5-CSRT task and performed at a level above chance (1 in 5 = 20% choice accuracy) in post-mTBI probe trials. Choice accuracy decreased (β = −1.84%/s, p < 2×10^-16^)^a^ and % omissions increased (β = 4.79%/s, p < 2×10^-16^)^b^ with decreasing stimulus duration, confirming that attention was challenged as the stimulus duration was decreased (**Figure 2**). Confirming the occurrence of head impact, mTBI mice experienced a short period of apnea immediately post-impact (apnea duration following the first mTBI: mean = 8.7 s, SD = 8.2 s; data available in a subset of 26 mice) whereas all sham mice experienced no apnea. Contrary to expectation, no effect of mTBI was observed on any of the 5-CSRT task measures at any of the timepoints or stimulus durations.

**Figure 2.**
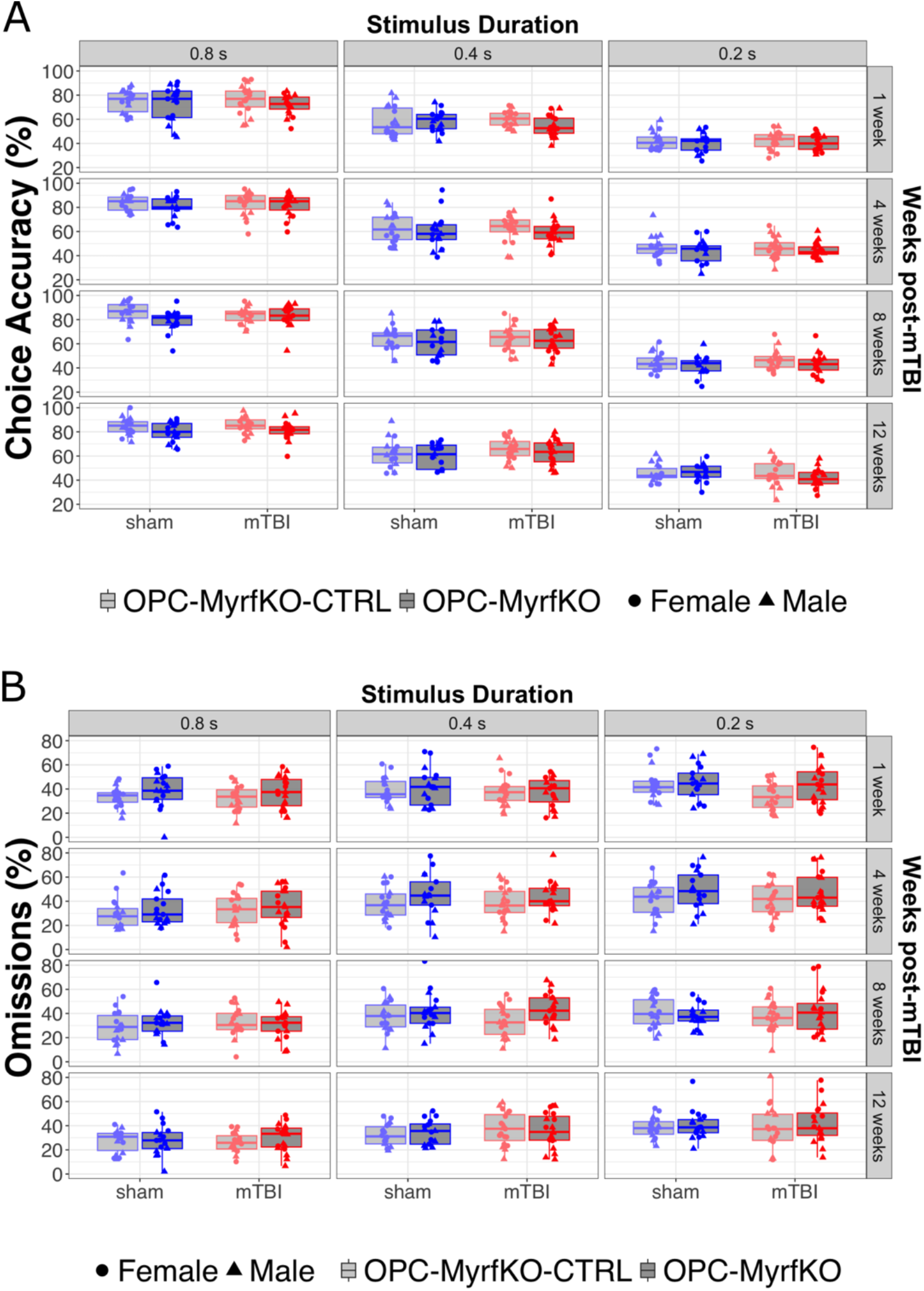
Choice accuracy (A) and omissions (B) on the 5-CSRT task in post-mTBI probe trials. Choice accuracy decreased and omissions increased with decreasing stimulus duration, confirming that attention was challenged with the shorter stimulus durations. OPC-MyrfKO mice had lower choice accuracy (β = −2.85%, p = 0.03) and more omissions (β = 4.09%, p = 0.02) than OPC-MyrfKO-CTRL mice across injury groups, stimulus durations and time points. No effect of mTBI was observed in these measures. For improved visualization, 1 data point in (A) and 3 data points in (B) are outside the y axis limits and not shown. All data points were included in the statistical models.

OPC-MyrfKO mice showed lower choice accuracy (β = −2.85%, p = 0.03)^c^ and omitted more trials (β = 4.09%, p = 0.02)^d^ than OPC-MyrfKO-CTRL mice across injury groups, probe trial stimulus durations and timepoints (**Figure 2**). Omissions were higher in OPC-MyrfKO compared to OPC-MyrfKO-CTRL mice during the inter-probe trial rebaselining sessions (2-s stimulus duration) as well (β = 1.37%, p = 0.05)^e^, but choice accuracy was similar between groups in these sessions (**Figure 2-1**). OPC-MyrfKO mice were slower to collect the reward (β = 0.09 s, p = 0.02)^f^ compared to OPC-MyrfKO-CTRL mice (**Figure 2-2**). A table of all statistical results is provided in the supplemental material (**Table 2-1**).

## 4 Discussion

Difficulty with focus and attention is common after mTBI in humans (Stojanovski et al., 2021) and has been reported in several rodent model studies (Mychasiuk et al., 2015; Xu et al., 2021; Zhang et al., 2021; Vonder Haar et al., 2022). Given the role for myelin in supporting and tuning neural circuit function (Noori et al., 2020), we aimed to test whether myelin plasticity was required for recovery of attention capabilities following mTBI. Although we did not observe impaired attention following mTBI in this study, precluding the ability to test the planned hypothesis, the intervention, halting adult oligodendrogenesis, did produce broad impairment on task performance, providing some insight into the potential role of new myelin in attention and possibly motivation and complex task performance.

The lack of an effect of mTBI on attention measures in the 5-CSRT task suggests that the model used in the current study may be too mild or may not have the injury characteristics required to elicit measurable impairment in attention. A similar mouse model that delivered one mTBI rather than three impacts was previously characterized with transient behavioural impairment observed three days after injury specific to the Y-maze test sensitive to working memory and attention (Gazdzinski et al., 2020). The same single impact model demonstrated persistent structural differences and neuroinflammation in white matter (Gazdzinski et al., 2020). That said, it is not uncommon for animal models of mTBI (Shultz et al., 2012; To and Nasrallah, 2021) as well as human neuroimaging studies (Manning et al., 2017; Churchill et al., 2019) to demonstrate ongoing differences in the brain following mTBI with no detectable behavioural differences or symptoms. Though previous studies have reported attentional impairment post-mTBI using the 5-CSRT task, they, in general, involved a higher number of impacts (>=5), incorporated a higher degree of rotational acceleration, were conducted in rats, or trained the animals after the injury, in which case task performance may be confounded by learning challenges post-mTBI (Mychasiuk et al., 2015; Xu et al., 2021; Vonder Haar et al., 2022).

Despite the absence of an attentional deficit following mTBI, the results suggest a role for oligodendrogenesis in attention and task engagement. Choice accuracy was high with the 0.8 s stimulus duration and decreased in all groups with decreasing stimulus duration, confirming that the mice learned the task well and that attention was being challenged with the shorter stimulus durations. Mice with halted oligodendrogenesis had lower choice accuracy and more omissions over all injury groups, time points and probe-trial stimulus durations. These mice were also slower to collect the reward following correct touches. These mice had similar choice accuracy as controls with the longer 2-s stimulus duration presented during the inter-probe trial rebaselining sessions, supporting the interpretation that attention is impaired when adult oligodendrogenesis is blocked. The higher omission rate and longer reward collection latency of these mice across injury groups, stimulus durations, and time points suggests that these mice may also be less motivated to perform the task.

Neuronal activity promotes oligodendrogenesis and adaptive myelination (Gibson et al., 2014) and oligodendrocytes provide support to axons through functions unrelated to myelin (Duncan et al., 2021; Krämer-Albers and Werner, 2023; Simons et al., 2024). These functions allow oligodendrocytes to contribute to experience dependent fine-tuning of neural circuits important for learning and memory (McKenzie et al., 2014; Xiao et al., 2016; Bacmeister et al., 2020; Pan et al., 2020; Steadman et al., 2020; Munyeshyaka and Fields, 2022; Shimizu et al., 2023; Yalçın et al., 2024). Reduced myelin turnover and remodeling reduces circuit plasticity, potentially impairing network function required for cognition. Altered patterns of neural activity and myelination have been reported in attention deficit and hyperactivity disorder (Lin et al., 2024; Gao et al., 2025; Gholamipourbarogh et al., 2025; Hu et al., 2025; Mizuno et al., 2025) and reduced neural activity and myelin content have been observed in several conditions associated with anhedonia and lack of motivation, including social deprivation (Liu et al., 2012; Hong et al., 2023), chronic stress (Kokkosis et al., 2022), and major depressive disorder (Sacchet and Gotlib, 2017). Relevant to the results presented here, using similar mice, a recent study by Yalcin et al. (Yalçın et al., 2024) demonstrated that active oligodendrogenesis is required for opioid-induced reward learning using the conditioned place preference (CPP) test. Interestingly, this did not translate to general reward behaviour such that the mice displayed normal food-induced CPP, naloxone-induced aversion learning, and sucrose preference. While food reward is a component of the 5-CSRT paradigm, the task is more complex than Pavlovian conditioning and therefore likely relies on coordinated activity across a distinct neural network. The details of this network and the mechanisms underlying the role of oligodendrogenesis in maintaining its function remain open questions.

There are limitations that should be considered while interpreting the results from this study. First, a comprehensive characterization of brain pathology following mTBI was not included, therefore the severity of the injury is assumed based on previous work. Several studies using similar models, including some involving a single impact, have reported pathological changes in the white matter assessed with histopathology and/or magnetic resonance imaging that persisted for several weeks to months post-injury (Yu et al., 2017; Lyons et al., 2018; Gazdzinski et al., 2020). Second, an assessment of the Myrf-KO on oligodendrocyte density or myelin was not included in this study. The persistence of the oligodendrogenesis blockade is assumed based on several previous studies using the same or similar model that have shown substantially reduced generation of new oligodendrocytes in these mice for at least 2-3 months post-tamoxifen administration (McKenzie et al., 2014; Pan et al., 2020; He et al., 2024; Kaller et al., 2024; Xin et al., 2024; Gazdzinski et al., 2025).

In conclusion, difficulty with attention is common after mTBI and can affect many aspects of cognition and quality of life. Identification of therapeutic targets is urgently needed and given myelin’s role in neural circuit function and its vulnerability to injury, myelin plasticity remains a potential candidate. While we did not observe impaired attention on the 5-CSRT task following mTBI in this study, mice unable to generate new myelin showed reduced task accuracy, possibly influenced by motivational processes. These findings suggest that adult oligodendrogenesis supports attention and complex behaviour. Further research may aim to optimize the injury model to better recapitulate the cognitive impairments observed in humans, including attention, or investigate the role of oligodendrogenesis in motivation with tests better suited to specifically measure this behaviour.

## Supporting information

Statistics table

## 5 Acknowledgements

The Myrf^*flox/flox*^ mice were a kind gift from Paul Frankland’s lab at the Hospital for Sick Children. The authors thank Xinyi Lin and Nicholas Wilson for their assistance with the 5-CSRT training.

## References

Armstrong RC, Mierzwa AJ, Marion CM, Sullivan GM (2016) White matter involvement after TBI: Clues to axon and myelin repair capacity. Exp Neurol.

Bacmeister CM, Barr HJ, McClain CR, Thornton MA, Nettles D, Welle CG, Hughes EG (2020) Motor learning promotes remyelination via new and surviving oligodendrocytes. Nat Neurosci 23:819–831.

Cancelliere C, Verville L, Stubbs JL, Yu H, Hincapié CA, Cassidy JD, Wong JJ, Shearer HM, Connell G, Southerst D, Howitt S, Guist B, Silverberg ND (2023) Post-Concussion Symptoms and Disability in Adults With Mild Traumatic Brain Injury: A Systematic Review and Meta-Analysis. J Neurotrauma.

Churchill NW, Hutchison MG, Graham SJ, Schweizer TA (2019) Mapping brain recovery after concussion: From acute injury to 1 year after medical clearance. Neurology 93:E1980–E1992.

Collins LK, Ofa SA, Miskimin C, Mulcahey M (2023) Cognitive Deficits Following Concussion: A Systematic Review. Journal of Orthopaedic Experience & Innovation 4.

Daugherty J, Depadilla L, Sarmiento K, Breiding MJ (2020) Self-Reported Lifetime Concussion among Adults: Comparison of 3 Different Survey Questions. Journal of Head Trauma Rehabilitation 35:E136–E143.

Duncan GJ, Manesh SB, Hilton BJ, Assinck P, Liu J, Moulson A, Plemel JR, Tetzlaff W (2018) Locomotor recovery following contusive spinal cord injury does not require oligodendrocyte remyelination. Nat Commun 9.

Duncan GJ, Simkins TJ, Emery B (2021) Neuron-Oligodendrocyte Interactions in the Structure and Integrity of Axons. Front Cell Dev Biol.

Fizet J, Cassel JC, Kelche C, Meunier H (2016) A review of the 5-Choice Serial Reaction Time (5-CSRT) task in different vertebrate models. Neurosci Biobehav Rev.

Gao Z, Duberg K, Warren SL, Zheng L, Hinshaw SP, Menon V, Cai W (2025) Reduced temporal and spatial stability of neural activity patterns predict cognitive control deficits in children with ADHD. Nature Communications 16.

Gazdzinski L, Mak J, Gumarathas K, Mellerup M, Collin A, Sled JG, Nieman BJ, Wheeler AL (2025) Oligodendrogenesis inhibition in the juvenile and adolescent periods differentially alters myelin in mice. Dev Neurosci 1–16.

Gazdzinski LM, Mellerup M, Wang T, Adel SAA, Lerch JP, Sled JG, Nieman BJ, Wheeler AL (2020) White Matter Changes Caused by Mild Traumatic Brain Injury in Mice Evaluated Using Neurite Orientation Dispersion and Density Imaging. J Neurotrauma 37:1818–1828.

Gholamipourbarogh N, Roessner V, Bluschke A, Beste C (2025) Novel Neural Activity Profiles Underlying Inhibitory Control Deficits of Clinical Relevance in Attention-Deficit/Hyperactivity Disorder: Insights From Electroencephalography Tensor Decomposition. Biol Psychiatry Cogn Neurosci Neuroimaging.

Gibson EM, Purger D, Mount CW, Goldstein AK, Lin GL, Wood LS, Inema I, Miller SE, Bieri G, Zuchero JB, Barres BA, Woo PJ, Vogel H, Monje M (2014) Neuronal activity promotes oligodendrogenesis and adaptive myelination in the mammalian brain. Science (1979) 344:1252304.

Glaser J, Jaeckle S, Beblo T, Mueller G, Eidenmueller AM, Schulz P, Schmehl I, Rogge W, Hollander K, Toepper M, Gonschorek AS (2024) The effect of repeated concussions on clinical and neurocognitive symptom severity in different contact sports. Scand J Med Sci Sports 34.

He Yuehua, Xu Z, He Yongxiang, Liu J, Li J, Wang S, Xiao L (2024) Preventing production of new oligodendrocytes impairs remyelination and sustains behavioural deficits after demyelination. Biochem Biophys Res Commun 733.

Hong H et al. (2023) Differential effects of social isolation on oligodendrocyte development in different brain regions: insights from a canine model. Front Cell Neurosci 17.

Hu R, Du W, Tan F, Wu Y, Yang C, Wang W, Chen W, Miao Y (2025) Dynamic alterations in spontaneous neural activity in patients with attention-deficit/hyperactivity disorder: A resting-state fMRI study. Brain Res Bull 222.

Johnson VE, Stewart W, Smith DH (2013) Axonal pathology in traumatic brain injury. Exp Neurol.

Kaller MS, Lazari A, Feng Y, van der Toorn A, Rühling S, Thomas CW, Shimizu T, Bannerman D, Vyazovskiy V, Richardson WD, Sampaio-Baptista C, Johansen-Berg H (2024) Ablation of oligodendrogenesis in adult mice alters brain microstructure and activity independently of behavioral deficits. Glia 72:1728–1745.

Kokkosis AG, Madeira MM, Mullahy MR, Tsirka SE (2022) Chronic stress disrupts the homeostasis and progeny progression of oligodendroglial lineage cells, associating immune oligodendrocytes with prefrontal cortex hypomyelination. Mol Psychiatry 27:2833–2848.

Krämer-Albers EM, Werner HB (2023) Mechanisms of axonal support by oligodendrocyte-derived extracellular vesicles. Nat Rev Neurosci.

Lin L, Chen Y, Dai Y, Yan Z, Zou M, Zhou Q, Qian L, Cui W, Liu M, Zhang H, Yang Z, Su S (2024) Quantification of myelination in children with attention-deficit/hyperactivity disorder: a comparative assessment with synthetic MRI and DTI. Eur Child Adolesc Psychiatry 33:1935–1944.

Liu J, Dietz K, Deloyht JM, Pedre X, Kelkar D, Kaur J, Vialou V, Lobo MK, Dietz DM, Nestler EJ, Dupree J, Casaccia P (2012) Impaired adult myelination in the prefrontal cortex of socially isolated mice. Nat Neurosci 15:1621–1623.

Lotocki G, de Rivero Vaccari JP, Alonso O, Molano JS, Nixon R, Safavi P, Dietrich WD, Bramlett HM (2011) Oligodendrocyte vulnerability following traumatic brain injury in rats. Neurosci Lett 499:143–148.

Lyons DN, Vekaria H, Macheda T, Bakshi V, Powell DK, Gold BT, Lin AL, Sullivan PG, Bachstetter AD (2018) A Mild Traumatic Brain Injury in Mice Produces Lasting Deficits in Brain Metabolism. J Neurotrauma 35:2435–2447.

Manning KY, Schranz A, Bartha R, Dekaban GA, Barreira C, Brown A, Fischer L, Asem K, Doherty TJ, Fraser DD, Holmes J, Menon RS (2017) Multiparametric MRI changes persist beyond recovery in concussed adolescent hockey players. Neurology 89:2157–2166.

Mar AC, Horner AE, Nilsson SRO, Alsiö J, Kent BA, Kim CH, Holmes A, Saksida LM, Bussey TJ (2013) The touchscreen operant platform for assessing executive function in rats and mice. Nat Protoc 8:1985–2005.

McKenzie IA, Ohayon D, Li H, de Faria JP, Emery B, Tohyama K, Richardson WD (2014) Motor skill learning requires active central myelination. Science (1979) 346:318–322.

Mierzwa AJ, Marion CM, Sullivan GM, Mcdaniel DP, Armstrong RC (2015) Components of Myelin Damage and Repair in the Progression of White Matter Pathology After Mild Traumatic Brain Injury.

Mizuno Y, Yamashita M, Shou Q, Hamatani S, Cai W (2025) A brief review of MRI studies in patients with attention-deficit/hyperactivity disorder and future perspectives. Brain Dev.

Munyeshyaka M, Fields RD (2022) Oligodendroglia are emerging players in several forms of learning and memory. Commun Biol.

Mychasiuk R, Hehar H, Esser MJ (2015) A mild traumatic brain injury (mTBI) induces secondary attention-deficit hyperactivity disorder-like symptomology in young rats. Behavioural Brain Research 286:285–292.

Nonaka M, Taylor WW, Bukalo O, Tucker LB, Fu AH, Kim Y, Mccabe JT, Holmes A (2021) Behavioral and Myelin-Related Abnormalities after Blast-Induced Mild Traumatic Brain Injury in Mice. J Neurotrauma 38:1551–1571.

Noori R, Park D, Griffiths JD, Bells S, Frankland PW, Mabbott D, Lefebvre J (2020) Activity-dependent myelination: A glial mechanism of oscillatory self-organization in large-scale brain networks. Proc Natl Acad Sci U S A 117:13227–13237.

Pan S, Mayoral SR, Choi HS, Chan JR, Kheirbek MA (2020) Preservation of a remote fear memory requires new myelin formation. Nat Neurosci 23:487–499.

Povlishock JT (1993) Pathobiology of Traumatically Induced Axonal Injury in Animals and Man. Ann Emerg Med 22.

R Core Team (2022) R: A Language and Environment for Statistical Computing.

Sacchet MD, Gotlib IH (2017) Myelination of the brain in major depressive disorder: An in vivo quantitative magnetic resonance imaging study. Sci Rep 7.

Shi H, Hu X, Leak RK, Shi Y, An C, Suenaga J, Chen J, Gao Y (2016) Demyelination as a Rational Therapeutic Target for Ischemic or Traumatic Brain Injury HHS Public Access. Exp Neurol.

Shimizu T, Nayar SG, Swire M, Jiang Y, Grist M, Kaller M, Sampaio Baptista C, Bannerman DM, Johansen-Berg H, Ogasawara K, Tohyama K, Li H, Richardson WD (2023) Oligodendrocyte dynamics dictate cognitive performance outcomes of working memory training in mice. Nat Commun 14.

Shultz SR, MacFabe DF, Foley KA, Taylor R, Cain DP (2012) Sub-concussive brain injury in the Long-Evans rat induces acute neuroinflammation in the absence of behavioral impairments. Behavioural Brain Research 229:145–152.

Simons M, Gibson EM, Nave K-A (2024) Oligodendrocytes: Myelination and axonal support. Cold Spring Harb Perspect Biol 16.

Smirl JD, Jones KE, Copeland P, Khatra O, Taylor EH, Van Donkelaar P (2019) Characterizing symptoms of traumatic brain injury in survivors of intimate partner violence. Brain Inj 33:1529–1538.

Steadman PE, Xia F, Ahmed M, Mocle AJ, Penning ARA, Geraghty AC, Steenland HW, Monje M, Josselyn SA, Frankland PW (2020) Disruption of Oligodendrogenesis Impairs Memory Consolidation in Adult Mice. Neuron 105:150–164.e6.

Stojanovski S, Scratch SE, Dunkley BT, Schachar R, Wheeler AL (2021) A Systematic Scoping Review of New Attention Problems Following Traumatic Brain Injury in Children. Front Neurol.

Sullivan GM, Mierzwa AJ, Kijpaisalratana N, Tang H, Wang Y, Song SK, Selwyn R, Armstrong RC (2013) Oligodendrocyte lineage and subventricular zone response to traumatic axonal injury in the corpus callosum. J Neuropathol Exp Neurol 72:1106–1125.

To XV, Nasrallah FA (2021) A roadmap of brain recovery in a mouse model of concussion: insights from neuroimaging. Acta Neuropathol Commun 9.

Vonder Haar C, Wampler SK, Bhatia HS, Ozga JE, Toegel C, Lake AD, Iames CW, Cabral CE, Martens KM (2022) Repeat Closed-Head Injury in Male Rats Impairs Attention but Causes Heterogeneous Outcomes in Multiple Measures of Impulsivity and Glial Pathology. Front Behav Neurosci 16.

Xiao L, Ohayon D, Mckenzie IA, Sinclair-Wilson A, Wright JL, Fudge AD, Emery B, Li H, Richardson WD (2016) Rapid production of new oligodendrocytes is required in the earliest stages of motor-skill learning. Nat Neurosci 19:1210–1217.

Xin W, Kaneko M, Roth RH, Zhang A, Nocera S, Ding JB, Stryker MP, Chan JR (2024) Oligodendrocytes and myelin limit neuronal plasticity in visual cortex. Nature.

Xu X et al. (2021) Repetitive mild traumatic brain injury in mice triggers a slowly developing cascade of long-term and persistent behavioral deficits and pathological changes. Acta Neuropathol Commun 9.

Yalçin B, Pomrenze MB, Malacon K, Drexler R, Rogers AE, Shamardani K, Chau IJ, Taylor KR, Ni L, Contreras-Esquivel D, Malenka RC, Monje M (2024) Myelin plasticity in the ventral tegmental area is required for opioid reward. Nature 630:677–685.

Yates LC, Yates E, Li X, Lu Y, Yakoub K, Davies D, Belli A, Sawlani V (2025) Developing a multivariate model for the prediction of concussion recovery in sportspeople: a machine learning approach. BMJ Open Sport Exerc Med 11.

Yu F, Shukla DK, Armstrong RC, Marion CM, Radomski KL, Selwyn RG, Dardzinski BJ (2017) Repetitive Model of Mild Traumatic Brain Injury Produces Cortical Abnormalities Detectable by Magnetic Resonance Diffusion Imaging, Histopathology, and Behavior. J Neurotrauma 34:1364–1381.

Zhang L, Levenson CW, Salazar VC, McCarthy DM, Biederman J, Zafonte R, Bhide PG (2021) Repetitive Mild Traumatic Brain Injury in a Perinatal Nicotine Exposure Mouse Model of Attention Deficit Hyperactivity Disorder. Dev Neurosci 43:63–72.

