## Supplementary figures and images for "Inhibited oligodendrogenesis, but not repeated mild traumatic brain injury, impairs attention in adult mice"

### Supplemental Figure 2-1

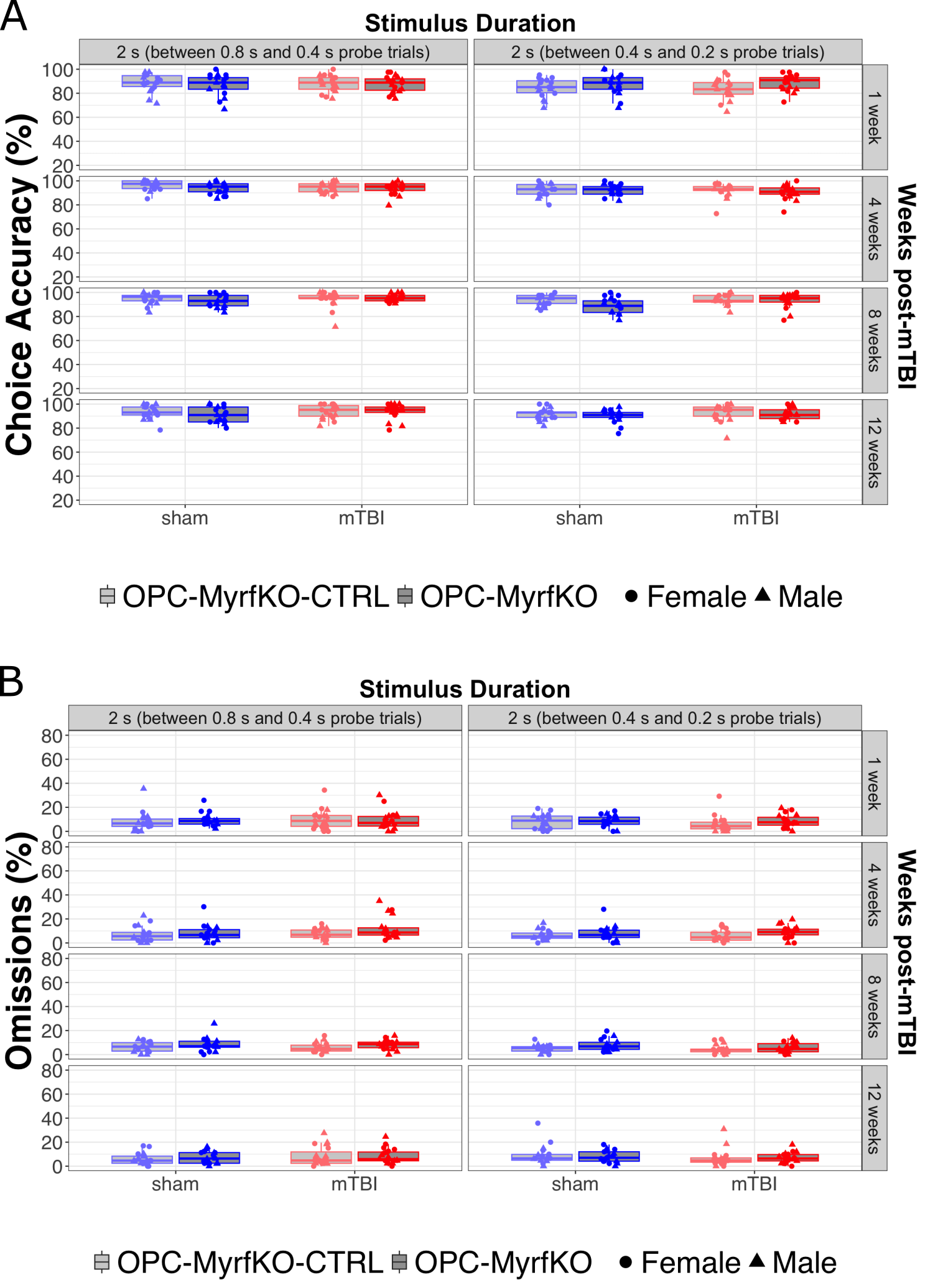

### Supplemental Figure 2-2

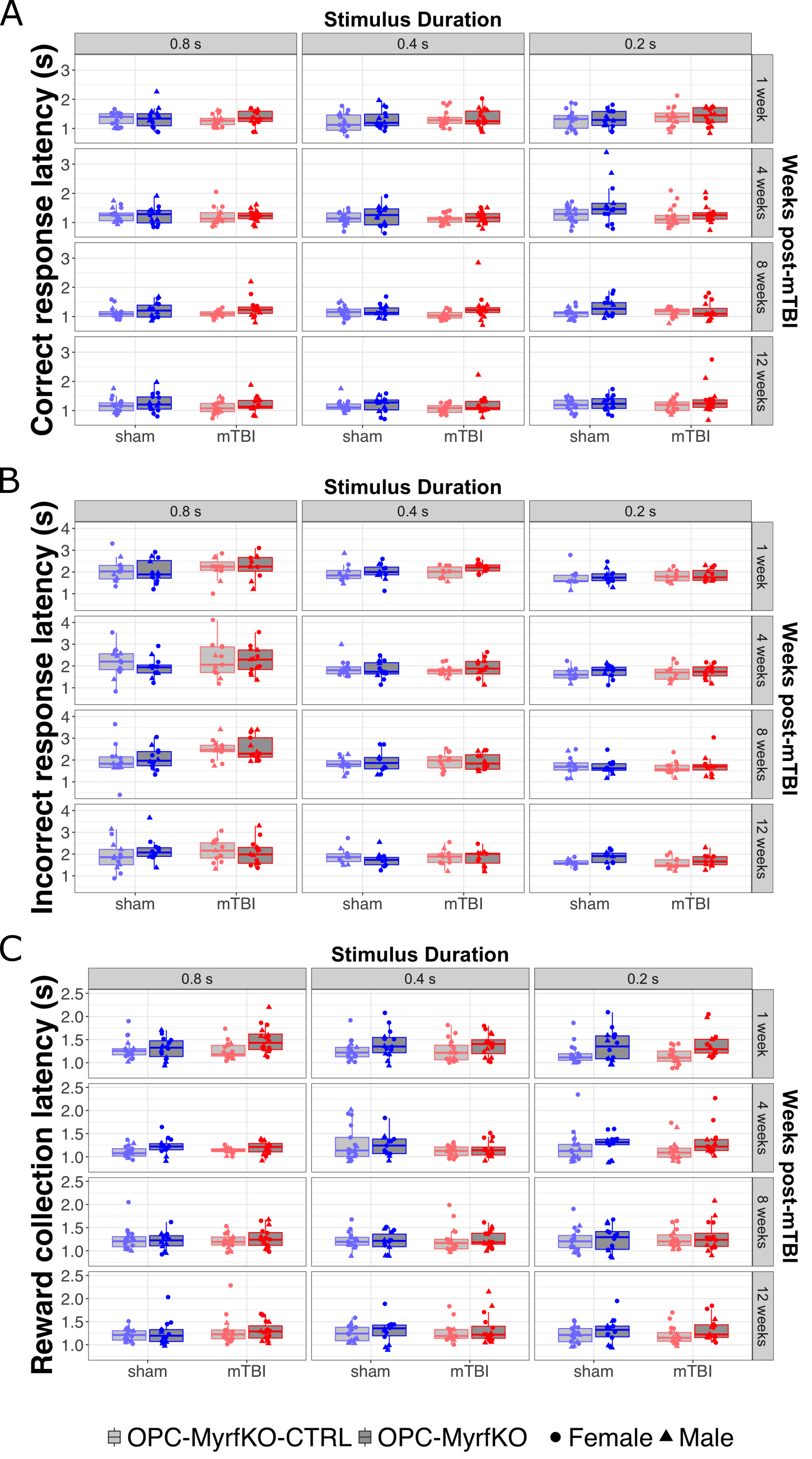
