## Supplemental methods for "Inhibited oligodendrogenesis, but not repeated mild traumatic brain injury, impairs attention in adult mice"

**Operant Chamber Habituation and Pre-Training Stages**

1. **Habituation**

- Mice were placed in the operant chambers for 30 min during which milkshake was delivered in the reward port. No stimulus was presented on the screen.
- Mice progressed to the next stage after completing two habituation sessions and consuming all of the milkshake delivered.

2. **Initial Touch (Touch Screen)**

- A stimulus window was illuminated for 30 s, changing location every 30 s, until the screen was touched.
- Milkshake reward was delivered for any screen touch, regardless of location.
- A 5-s inter-trial interval (ITI) began once the mouse collected the reward and exited the reward port, after which a new stimulus was presented on the screen.
- Mice progressed to the next stage once they completed 30 trials within 60 min on two consecutive days.

3. **Must Touch Stimulus**

- A stimulus window was illuminated until it was touched, after which milkshake reward was delivered. No reward was delivered for touching the screen in a location that was not illuminated.
- Mice progressed to the next stage once they completed 30 trials within 60 min on two consecutive days.

4. **Must Initiate**

- As Stage 3, but the mouse was required to re-enter and exit the reward port after the ITI to initiate a new trial. This allowed the mice to self-pace the session.
- Mice progressed to the next stage once they completed 30 trials within 60 min on three consecutive days.

5. **Introduction to Timeouts**

- As Stage 4, but the mouse was ‘punished’ for incorrect touches (i.e. touching the screen in a location other than the illuminated window) with a 5-s timeout period during which a buzzer sounded and the overhead chamber light was illuminated. The mouse was able to initiate a new trial following a 5-s ITI that occurred after the timeout period.
- Mice progressed to task-specific training once they completed 18 correct trials within 35 mins on three of four consecutive days. The shorter session duration and lower success criteria compared to previous stages were chosen to minimize the stress imposed by the timeout periods while maintaining an equivalent correct trial rate requirement.

**Additional notes**

- Mice were trained 6 days/week (Monday-Saturday).
- If mice were unable to reach criteria on a given stage for a week, they were moved back to the previous stage until able to reach criteria again before progressing.
